# A genetic variant at coronary artery disease and ischemic stroke locus 1p32.2 regulates endothelial responses to hemodynamics

**DOI:** 10.1101/377549

**Authors:** Matthew D. Krause, Ru-Ting Huang, David Wu, Tzu-Pin Shentu, Devin L. Harrison, Michael B. Whalen, Lindsey K. Stolze, Anna Di Rienzo, Ivan P. Moskowitz, Mete Civelek, Casey E. Romanoski, Yun Fang

**Affiliations:** Department of Medicine, The University of Chicago; Department of Cellular and Molecular Medicine, The University of Arizona; Departments of Human Genetics; Departments of Pediatrics and Pathology, The University of Chicago; Department of Biomedical Engineering, The University of Virginia

**Author notes:** Correspondence to Yun Fang, PhD.

## Abstract

Biomechanical cues dynamically control major cellular processes but whether genetic variants actively participate in mechano-sensing mechanisms remains unexplored. Vascular homeostasis is tightly regulated by hemodynamics. Exposure to disturbed blood flow at arterial sites of branching and bifurcation causes constitutive activation of vascular endothelium contributing to atherosclerosis, the major cause of coronary artery disease (CAD) and ischemic stroke (IS). Conversely, unidirectional flow promotes quiescent endothelium. Genome-wide association studies have identified chromosome 1p32.2 as one of the most strongly associated loci with CAD/IS; however, the causal mechanism related to this locus remains unknown. Employing statistical analyses, ATAC-seq, and H3K27ac/H3K4me2 ChIP-Seq in human aortic endothelium (HAEC), our results demonstrate that rs17114036, a common noncoding polymorphism at the 1p32.2, is located in an endothelial enhancer dynamically regulated by hemodynamics. CRISPR/Cas9-based genome editing shows that rs17114036-containing region promotes endothelial quiescence under unidirectional flow by regulating phospholipid phosphatase 3 (PLPP3). Chromatin accessibility quantitative trait locus mapping using HAECs from 56 donors, allelic imbalance assay from 7 donors, and luciferase assays further demonstrate that CAD/IS protective allele at rs17114036 in PLPP3 intron 5 confers an increased endothelial enhancer activity. ChIPPCR and luciferase assays show that CAD/IS protective allele at rs17114036 creates a binding site for transcription factor Krüppel-like factor 2, which increases the enhancer activity under unidirectional flow. These results demonstrate for the first time that a human single-nucleotide polymorphism contributes to critical endothelial mechanotransduction mechanisms and suggest that human haplotypes and related *cis*regulatory elements provide a previously unappreciated layer of regulatory control in cellular mechano-sensing mechanisms.

**Significance Statement:** Biomechanical stimuli control major cellular functions and play critical roles in the pathogenesis of diverse human diseases. Although recent studies have implicated genetic variation in regulating key biological processes, whether human genetic variants contribute to the cellular mechano-sensing mechanisms remains unclear. This study provides the first line of evidence supporting an underappreciated role of genetic predisposition in cellular mechanotransduction mechanisms. Employing epigenomic profiling, genome-editing, and latest human genetics approaches, our data demonstrate that rs17114036, a common noncoding polymorphism implicated in coronary artery disease and ischemic stroke by genome-wide association studies, dynamically regulates endothelial responses to blood flow (hemodynamics) related to atherosclerosis via regulation of an intronic enhancer. The results provide new molecular insights linking disease-associated genetic variants to cellular mechanobiology.

## Introduction

Mechanical stimuli regulate major cellular functions and play critical roles in the pathogenesis of diverse human diseases (1). This is especially important in the vasculature, where endothelial cells are activated by local disturbed flow in arterial regions prone to atherosclerosis (2–6), the major cause of coronary artery disease (CAD) and ischemic stroke (IS). The role of biomechanical forces on the non-coding and regulatory regions of the human genome is unexplored. Recent studies demonstrated that the non-coding, non-transcribed human genome is enriched in *cis*regulatory elements (7). In particular, enhancers are distinct genomic regions that contain binding sites for sequence-specific transcription factors (8). Enhancers spatially and temporally control gene expression with cell type and cell state specific patterns (9). Notably, top-associated human disease-associated single nucleotide polymorphisms (SNPs) are frequently located within enhancers that explicitly activate genes in disease relevant cell types (10). The nature of mechano-sensitive enhancers and their biological roles in vascular functions have not been identified.

Atherosclerotic disease is the leading cause of morbidity and mortality worldwide. Genome-wide association studies (GWAS) identified chromosome 1p32.2 as one of the loci most strongly associated with susceptibility to CAD and IS (11–13). One candidate gene in this locus is PhosphoLipid PhosPhatase 3 (PLPP3, also known as PhosPhatidic-Acid-Phosphatase-type-2B/PPAP2B) which inhibits endothelial inflammation and promotes monolayer integrity by hydrolyzing lysophosphatidic acid (LPA) that activates endothelium (14, 15). Our recent study demonstrated that PLPP3 expression is significantly increased in vascular endothelium by unidirectional flow *in vitro* and *in vivo* (14). Moreover, expression quantitative trait locus (eQTL) mapping showed that CAD protective allele at 1p32.2 is associated with increased PLPP3 expression in an endothelium-specific manner (14). However, whether genetic variants and mechano-sensing mechanisms converge on PLPP3 expression is unclear. In addition, causal SNP(s) at locus 1p32.2 remain unknown.

Employing statistical analyses, ATAC-seq, H3K27ac ChIP-seq, H3K4me2 ChIP-seq, luciferase assays, and CRISPR/Cas9-based genome editing, we report that rs17114036-containing genomic region at 1p32.2 causatively promotes endothelial expression of phospholipid phosphatase 3 (PLPP3) and governs the athero-resistant endothelial phenotype under unidirectional flow by functioning as a mechano-sensitive endothelial enhancer. Using HAECs isolated from a cohort of human subjects, we performed transcriptome analyses and chromatin accessibility quantitative trait locus (caQTL) mapping showing nucleotide-specific epigenetic and transcriptomic effects of rs17114036 in humans. Allelic imbalance assays, ChIP-PCR, and luciferase assays collectively demonstrate that due to a single base pair change, the CAD/IS protective allele at rs17114036 confers increased activity of an endothelial intronic enhancer that is dynamically activated by unidirectional blood flow and transcription factor Krüppel-like factor 2 (KLF2). This is the first report elucidating underlying molecular mechanisms related to CAD/IS locus 1p32.2 and linking human disease-associated genetic variants to critical mechano-transduction mechanisms. The new molecular insights suggest that human genetic variants provide a novel layer of molecular control by which cells convert physical stimuli into biological signaling via tissue-specific enhancers.

## Results

### Bayesin Refinement and Conditional and Joint multiple-SNP analyses predict rs17114036 and rs2184104 are putative causal SNPs located in CAD/IS locus 1p32.2

Rs17114036 is the tag SNP used in most CAD/IS GWAS (11–13) and in our eQTL mapping (14); however, there are forty-four common SNPs in high linkage disequilibrium (LD) (r^2^>0.8) with rs17114036, and any of these SNPs could conceivably be a causal variant. To predict possible causal SNPs at the 1p32.2 locus, we conducted two statistical analyses. First, we employed a Bayesian statistical approach to assign posterior probabilities and credible sets of SNPs that refine the association signals of GWAS-detected loci (16). Second, we applied conditional and joint association analysis using summary-level statistics of GWAS data to predict causal variants (17). Using Bayes’ theorem in the cohort of forty-five SNPs at 1p32.2, we identified fifteen SNPs with >95% posterior probability to be causal (Fig. 1A and Supplementary Table 1). Using the approximate conditional and joint association analysis, we identified seven 1p32.2-associated SNPs to be possible causal (Fig. 1A and Supplementary Table 1). Only two SNPs, rs17114036 and rs2184104, were predicted to be causal by both methods.

**Figure 1.**
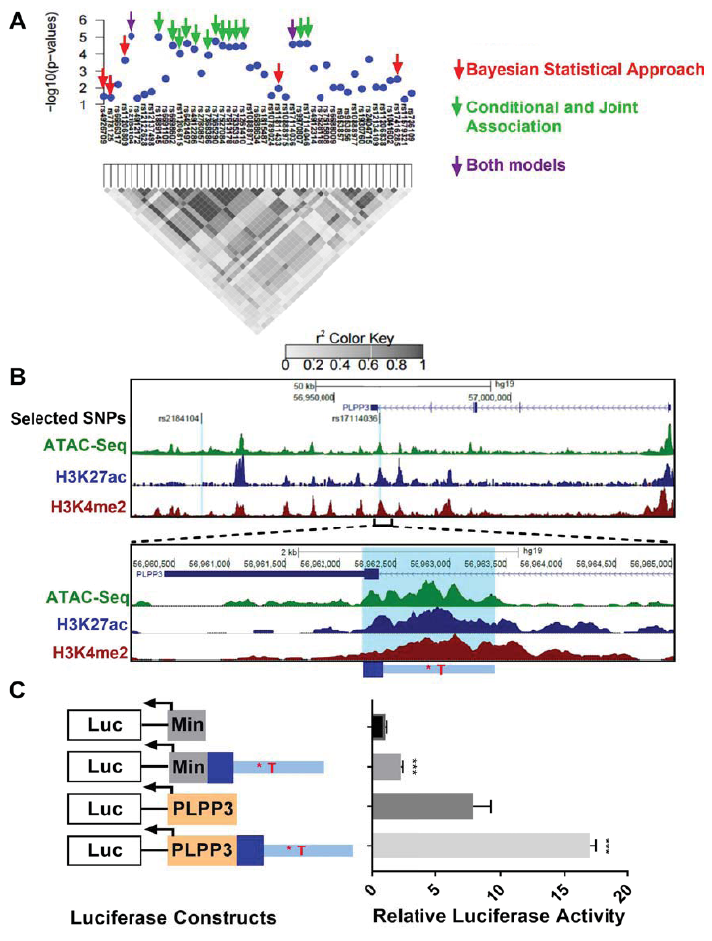
CAD-associated SNP rs17114036 is located in an enhancer element (chr1:56962213-56963412) in human aortic endothelial cells (HAEC). **(A)** In silico prediction of causal SNPs in the CAD locus 1p32.2. Diagrams demonstrate the association (-log10(P) and link)age disequilibrium (LD) pattern of total 45 common SNPs in the 1p32.2 locus. Red arrows indicate possible causal SNPs predicted by Bayesian Statistical Approach. Green Harrows indicate possible causal SNPs predicted by Approximate Conditional and Joint Association Analysis. Purple arrows indicate putative causal SNPs predicted by both statistical analyses. **(B)** Chromatin accessibility and canonical enhancer marks in chr1:56962213-56963412 region enclosing rs17114036 in HAEC. Assay of Transposase Accessible Chromatin with whole genome sequencing (ATAC-seq) and H3K27ac/H3K4me2 chromatin immuno-precipitation with whole genome sequencing (ChIP-seq) collectively identified an enhancer-like region in chr1:56962213-56963412 in HAEC. All sequencing experiments were performed in duplicate and the merged tracks are shown. **(C)** Enhancer activity of chr1:56962213-56963412 in vascular endothelium. DNA sequences of chr1:56962213-56963412 were cloned into luciferase reporters (firefly luciferase construct PGL4) that contain a minimal promoter or human PLPP3 promoter. Dual-luciferase reporter assays were conducted in HAEC 24 hours after the electroporation-based transfection (PRL-TK of Renilla luciferase as transfection controls) in HAEC, detecting increased firefly luciferase as the result of insertion of chr1:56962213-56963412.

### CAD/IS-associated SNP rs17114036 is located in an enhancer element (chr1:56497123-56497188) in human aortic endothelial cells

Both rs17114036 and rs2184104 are located in non-coding regions. To probe the regulatory functions of these two SNPs in vascular endothelium, we performed Assay of Transposase Accessible Chromatin with whole genome sequencing (ATAC-seq) as well as H3K27ac and H3K4me2 chromatin immuno-precipitation with whole genome sequencing (ChIP-seq) in human aortic endothelial cells (HAECs). ATAC-seq is a high-throughput, genome-wide method to define chromatin accessibility that correlates with precise measures of transcription factor binding (18). The combination of H3K27ac and H3K4me2 ChIP-seq marks were used to identify active enhancers. Since the human PLPP3 gene is expressed from the minus strand in the annotated human genome, we use alleles in the minus strand at rs17114036 and rs2184104 in this manuscript. It is important to note that since CAD/IS risk alleles at rs17114036 (T) and rs2184104 (A) are major alleles in all ethnic groups (70-99% frequency) (19), our experiments, unless specified otherwise, were conducted in HAEC lines from donors who carry major alleles at rs17114036 and rs2184104. As demonstrated in Fig. 1B, rs17114036 in the intron 5 of the PLPP3 resides in an enhancer-like element (chr1:56962213-56963412, UCSC VERSION hg19) identified by ATAC-seq and H3K27ac/H3K4me2 ChIP-seq in HAECs. Encyclopedia of DNA Elements (ENCODE) also reported a DNase hypersensitive site and a H3K27ac peak in a ~1kb region enclosing rs17114036 in human umbilical vein endothelial cells (HUVEC) (7) (Supplemental Fig. 1). Notably, this region does not exhibit enhancer-like marks in other ENCODE cell types, such as K562, GM12878, and NHEK cells (Supplemental Fig. 1). In contrast, the other putative causal SNP, rs2184104, is located ~120kb downstream of the PLPP3 transcription start site at a location that lacks enhancer-like features (Fig. 1B). ENCODE data are consistent with our findings, and also signify an inactive chromatin domain surrounding rs2184104 (Supplemental Fig 1). Supplemental Fig. 2 shows the ATAC-seq and H3K27ac/H3K4me2 tracks in HAECs at 1p32.2 locus. The enhancer activity of chr1:56496541-56497740 was experimentally demonstrated by a luciferase reporter assay (Fig. 1C). Plasmid transfection was first validated in HAECs using electroporation of pmaxGFP-expressing constructs (Supplemental Fig. 3). A 1200 bp DNA sequence corresponding to human chr1:56962213-56963412 was cloned upstream of firefly luciferase that was driven by a minimal promoter. Reporter assays demonstrated that insertion of this putative enhancer region with major allele T at rs17114036 significantly increased the luciferase activity in HAECs (Fig. 1C). We further cloned these putative enhancer elements in a luciferase vector that contains the human PLPP3 promoter. Endogenous human PLPP3 promoter led to a 7.9 fold higher luciferase activity in HAECs when compared with the vector with minimal promoter (Fig. 1C). Moreover, insertion of the putative enhancer elements resulted in a 2.14-fold increase in luciferase activity when compared with the vector with only PLPP3 promoter (Fig. 1C). Minimal enhancer activities were detected when the constructs were expressed in the non-endothelium cell line HEK 293 (Supplemental Fig. 4). ATAC-seq, H3K27ac/H3K4me2 ChIP-seq, and luciferase assays altogether demonstrate that chr1:56962213-56963412 functions as an enhancer in HAECs.

**Figure 2.**
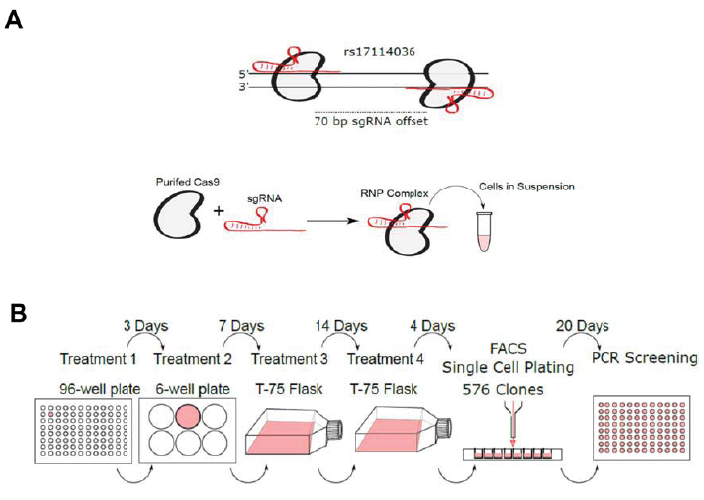
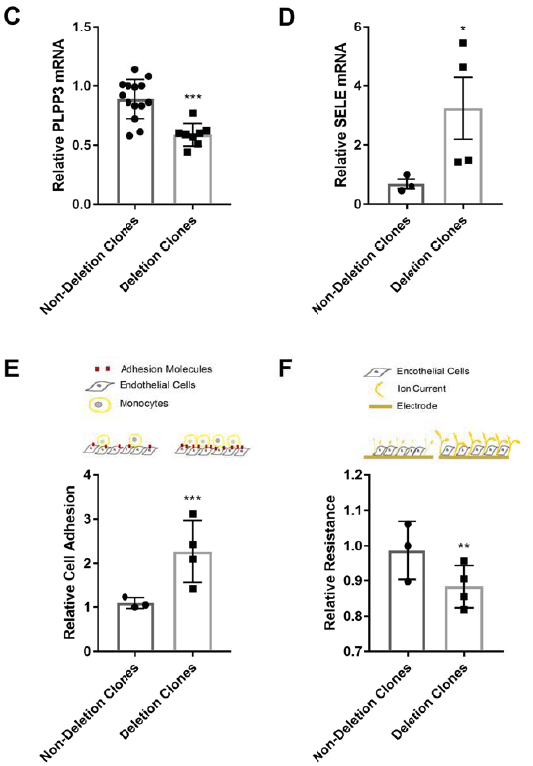
CRISPR/Cas9-mediated deletion of rs17114036-containing genomic locus reduces PLPP3 expression, promotes inflammation, and increases monolayer permeability in human aortic endothelium. (**A**) RNA ribonucleoprotein (RNP) complex thatcontains recombinant *S. pyogenes* Cas9 and two guide RNAs (gRNAs) flanking rs17114036.(**B**) Experimental overview of CRISPR/Cas9-mediated genomic deletion in HAEC. TeloHAEC were treated with RNP complex 4 times before single-cell sorting and isogenic clone selection. (**C1) Reduced PLPP3 expression, (**D**) Elevated E-selectin expression, (**E**) Increased leukocyte adhesion, and (**F**) Highermonolayer permeability in teloHAEC with genomic deletion at rs17114036-contaning region. n=3-8. **p<0.005; ***p<0.0005 as determinedby Student’s t-test. Data represent mean ± SEM.**

### CRISPR/Cas9-mediated deletion of rs17114036-containing genetic locus reduces PLPP3 expression in human aortic endothelium

To determine the causal role of rs17114036-containing genomic locus in regulating endothelial PLPP3 expression, the bacterial CRISPR-associated protein-9 nuclease (CRISPR/Cas9) system was used to selectively delete a ~66 bp genomic region (chr1:56962783-56962849) enclosing rs17114036 in HAECs. A pair of guide RNAs (Fig. 2A) was designed according to the method described by Ran et al (20). We pre-assembled Cas9-guide RNA ribonucleoprotein (RNP) complex by incubating guide RNAs with recombinant *S. pyogenes* Cas9 (21), followed by the delivery to cells using cationic liposome transfection reagents. To improve the efficiency of the CRISPR/Cas9-mediated deletion, cells were reverse-transfected by the RNP complex four times before flow cytometry assays to sort single cells (Fig. 2B). Immortalized HAECs (carrying major alleles at rs17114036) with high proliferating capacity were used for single cell clonal isolation that selects a genetically-identical cell line. Among 459 HAEC colonies we grew to confluency, PCR assays detected 17 lines with ~66 bp genetic deletion in PLPP3 intron 5 enclosing rs17114036 (Supplemental Fig. 5A). DNA deletion was further confirmed by TA-cloning and Sanger sequencing (Supplemental Fig 5B). Endothelial PLPP3 expression is significantly reduced in the genome-edited cells when compared to teloHAECs that underwent CRISPR/Cas9 treatment and single cell clonal isolation but showed no sign of deletion at chr1:56962783-56962849 (Fig 2C). In addition, deletion of this putative enhancer in PLPP3 intron 5 resulted in an increase of LPA-induced E-selectin expression (Fig 2D) and leukocyte adhesion (Fig 2E), in agreement with the anti-inflammatory/adhesive role of endothelial PLPP3 (14, 15). Moreover, trans-endothelial electrical resistance (TER) detected increased monolayer permeability in rs17114036-deleted HAECs (Fig 2F), consistent with PLPP3’s role in maintaining endothelial monolayer integrity (14, 15). These results demonstrate that deletion of the rs17114036-containing region in HAECs causatively reduces PLPP3 expression and promotes endothelial activation.

### Unidirectional flow increases the enhancer activity at chr1:56497123-56497188 in vascular endothelium

Given the critical role of hemodynamics in controlling endothelial PLPP3 transcription (14), we tested whether blood flow regulates the enhancer activity of chr1:56962213-56963412. ATAC-seq and H3K27ac ChIP-seq were conducted in HAECs subjected to “athero-protective” unidirectional flow representing wall shear stress in human distal internal carotid artery or “athero-susceptible” flow mimicking hemodynamics in human carotid sinus (22). ATAC-seq captured an increased open chromatin region at chr1:56962213-56963412 in HAECs under unidirectional flow compared with cells under disturbed flow (Fig 3A). H3K27ac ChIP-seq indicated an increased enhancer activity of chr1:56962213-56963412 in HAECs under unidirectional flow (Fig 3A). In addition, genetic deletion of the rs17114036-containing region by CRISPR/Cas9 significantly impaired unidirectional flow-induced PLPP3 expression in HAECs (Fig 3B). These results collectively demonstrate that enhancer activity of chr1:56962213-56963412 is dynamically activated by the athero-protective unidirectional flow to regulate endothelial PLPP3.

**Figure 3.**
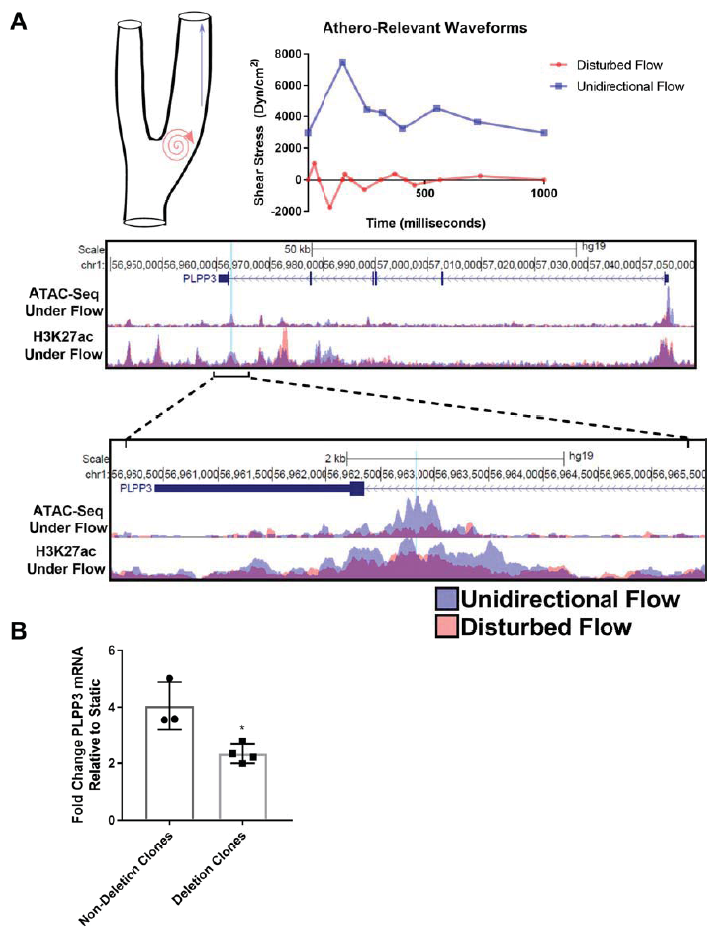
Unidirectional flow increasesenhancer activity at chr1:56962213-56963412 thattranscriptionally activates PLPP3 in human aortic endothelium. (A) Increased chromatin accessibility and H3K27ac mark at chr1:56962213-56963412 in HAEC subjected to 24-hr athero-pr tective unidirectional flow (UF) when compared with cells under 24-hr athero-susceptible disturbed flow (DF). The PLPP3 locus is shown and zoomed in to demonstrate details around the enhancer region of interest with rs17114036 highlighted with a vertical blue line. **(B)** Reduced UF-induced PLPP3 expression in HAEC with genomic deletion at rs17114036-containing genomic locus. Control and genome-edited (~66 bp deletion) HAEC were subjected to 24-hr unidirectional flow. Y-axis represents the fold change of PLPP3 mRNA quantities between the static conditions and unidirectional flow for each individual clone. Non-deletion clones n=3, deletion clones n=4. *p<0.05 as determined by Student’s t-test. Data represent mean ± SEM.

### CAD/IS protective allele C at rs1711403 confers a higher enhancer activity of chr1:56497123-56497188

GWAS have linked the minor allele C at rs17114036 at 1p32.2 to reduced CAD/IS susceptibility (11–13) and our eQTL mapping described increased PLPP3 expression in HAECs with minor allele C (14). We investigated the genotype-dependent effect of rs17114036 on the enhancer activity of chr1:56962213-56963412 by ATAC-seq and luciferase assays. In addition to HAEC lines carrying major (risk) allele T at rs17114036, we conducted ATAC-seq in HAECs isolated from donors who are heterozygous (T/C) (~20% of Europeans) at rs17114036, allowing us to perform chromatin accessibility quantitative trait locus (caQTL) mapping. caQTL was recently developed to detect between-individual signaling in *cis*-regulatory element as a function of genetic variants (23). ATAC-seq detected significantly increased numbers of reads corresponding to rs17114036-containing region in HAEC lines that contain one CAD protective allele (T/C) when compared with HAECs from donors homozygous of CAD risk allele (T/T) (Fig 4A), supporting increased chromatin accessibility associated with C allele at rs17114036. In addition, we conducted RNA-seq analysis in these cells demonstrating that there is a strong correlation between enhanced chromatin accessibility in rs17114036-containing region and increased mRNA levels of PLPP3 in HAECs (Fig 4B), further suggesting that chr1:56962213-56963412 functions as an enhancer in promoting endothelial PLPP3 transcription. Moreover, ATAC-seq experiments in HAEC lines heterozygous at rs17114036 further allow us to determine whether the chromosome with C at rs17114036 exhibits higher chromatin accessibility at chr1:56962213-56963412 when compared to the chromosome with T allele. This is achieved by the allelic imbalance (AI) analysis which assigns next generation sequencing reads overlapping heterozygous sites to one chromosome or the other for allele-specific signals (24). ATAC-seq detected reads enriched from the C-containing chromosome compared to that with T allele in HAECs heterozygous at rs1711403 (Fig. 4C), further supporting the increased chromosome accessibility associated with C allele at rs17114036. Lastly, luciferase assays were conducted to support the genotype-dependent enhancer activity of chr1:56962213-56963412. Replacement of T allele with C allele led to a much higher luciferase activity (~5.2 fold, C vs T) in endothelium (Fig. 4D). Taken together, these results demonstrate that CAD protective allele C at rs17114036 confers a higher enhancer activity of chr1:56962213-56963412 to promote PLPP3 expression in vascular endothelium.

**Figure 4.**
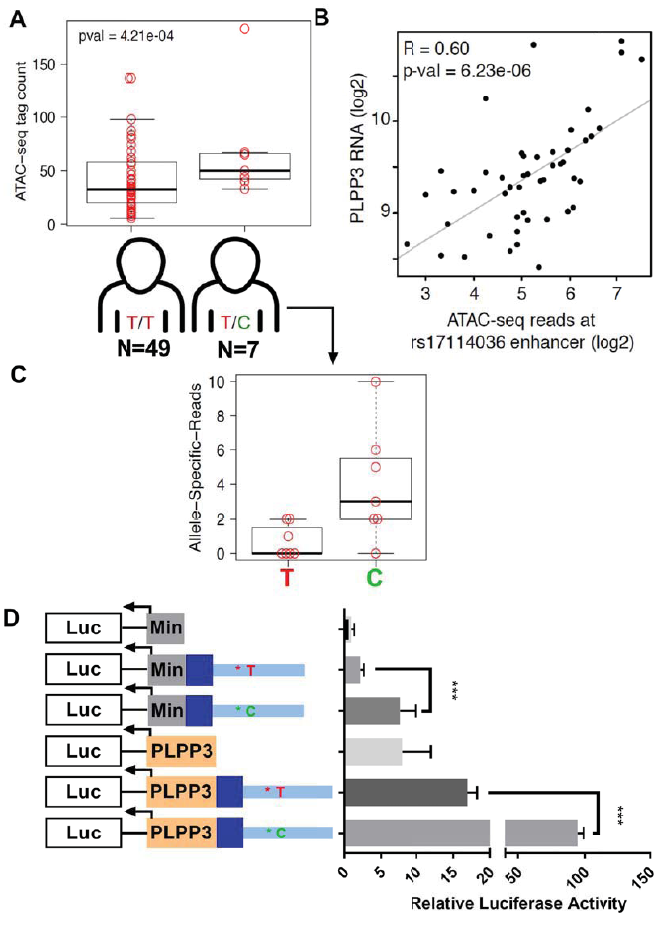
CAD protective allele C at rs17114036 confers higher enhancer activity of chr1:56962213-56963412. **(A)** Increased ATAC-seq reads in chr1:56962213-56963412 region from HAEC isolated from people heterogeneous (T/C) at rs17114036 when compared with HAEC from people heterozygous (T/T) at rs17114036. **(B)** A positive correlation (R=0.6, P-value=6.23e-06) between ATAC-seq reads at chr1:56962213-56963412 and PLPP3 mRNA detected by RNA-seq in 56 HAEC lines. **(C)** Increased ATAC-seq reads at rs17114036-containing genomic locus from C (rs17114036)-containing chromosome compared with T-containing chromosome in HAEC lines heterozygous (T/C) at rs17114036. **(D)** Increased enhancer activity of chr1:56962213-56963412 with C allele at rs17114036 when compared with T allele. Dual-luciferase reporter assays were conducted in teloHAEC. n=4-6. ***p<0.0005 as determined by Student’s t-test. Data represent mean ± SEM.

### CAD/IS protective C allele at rs1711403 promotes flow-induced, KLF2-mediated enhancer activity of chr1:56497123-56497188

We further examined whether the genetic variants at rs17114036 modulate the flow-induced enhancer activity of chr1:56962213-56963412. Luciferase assays detected an increase of the enhancer activity of chr1:56962213-56963412 (with protective C allele at rs17114036) in cells under 18-hr unidirectional flow when compared to disturbed flow (Fig 5A). In addition to the ATAC-seq experiments in HAEC homozygous at rs17114036 under flow (Fig. 3), we performed ATAC-seq analysis in four HAEC lines heterozygous at rs17114036 under 24-hr unidirectional flow, in order to perform open chromatin allelic imbalance (AI) analysis. Supplemental Fig. 6 demonstrates that in all four selected HAEC lines heterozygous at rs17114036, unidirectional flow increases ATAC-seq peaks in the proposed enhancer region in PLPP3 intron 5, in agreement with increased ATAC-seq reads in rs17114036-containging region (Fig. 5B). Moreover, allelic imbalance analysis showed an enrichment of ATAC-reads from the chromosome harboring the protective C allele (Fig. 5B). In contrast, ATAC-seq detected no allelic imbalance at rs6421497, a common SNP in high LD with rs17114036 (Supplemental Fig. 7). Indeed, the protective allele C at rs17114036 creates a CACC box that is a binding site for KLF2 which mediates the flow sensitivity of a cohort of endothelial genes including PLPP3 (14, 25-27). We then tested whether KLF2 dynamically activates this rs17114036-containing enhancer and if rs17114036 alleles impact the KLF2-mediated enhancer activity. First, the affinity of KLF2 to the rs17114036-containing locus was determined by KLF2 chromatin immunoprecipitation polymerase chain reaction (ChIPPCR) assays in HAECs carrying a protective allele at rs17114036, showing a physical binding of KLF2 to the CACC sites in the PLPP3 promoter and at rs17114036 (Fig. 5C). Enhancer activities of chr1:56962213-56963412 was further determined in HAECs as a function of KLF2 expression. Constructs of enhancer (chr1:56962213-56963412) and PLPP3 promoter were co-transfected with KLF2-overexpressing plasmids. Luciferase assays detected a 2.9 fold increase of luciferase activity in the T allele-containing construct as the result of KLF2 overexpression (Fig. 5D). Moreover, KLF2 overexpression led to a 4.7 fold increase of luciferase activity when T allele was substituted by the protective allele C at rs17114036 (Fig. 5D). Collectively, KLF2 ChIPPCR and luciferase assays demonstrate that CAD/IS protective allele C at rs17114036 confers a higher KLF2-dependent enhancer activity of chr1:56962213-56963412 in vascular endothelium.

**Figure 5.**
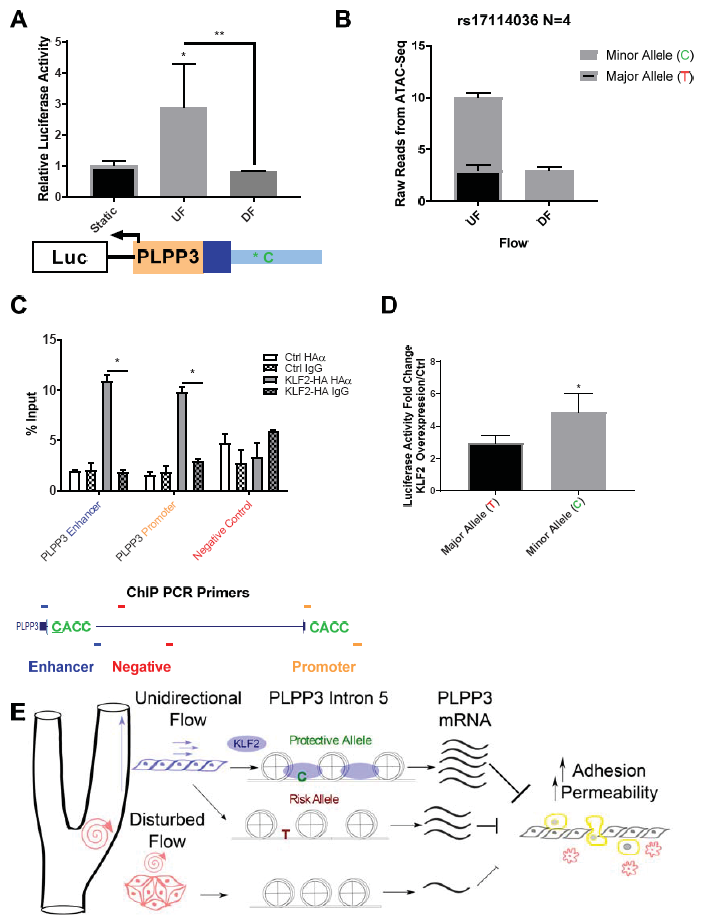
CAD protective C allele at rs17114036 promotes flow-induced, KLF2-mediated enhancer activity of chr1:56962213-56963412. **(A)** Increased enhancer activity of chr1:56497123-56497188 (with C allele at rs17114036) under unidirectional flow but not disturbed flow. n=4. **(B)** Increased ATAC-seq reads in rs17114036-containing region in HAEC under unidirectional flow (UF) when compared with disturbed flow (DF). ATAC-seq was conducted in 4 HAEC lines heterozygous at rs17114036 under 24-hr atherorelevant flow, detecting increased ATAC reads in cells under UF and higher reads from the C allele-containing chromosome compared with T allele. **(C)** KLF2 affinity to CACC sites in human PLPP3 promoter and intron 5. ChIP-quantitative polymerase chain reaction (ChIP-PCR) was performed with either a control IgG antibody or the antibody against HA followed by quantitative PCR using primers detecting CACC sites in PLPP3 promoter or rs17114036-enclsoing region from control HAEC (Ctrl) or HAEC transfected with KLF2 transcripts with HA tag. Primers that detect a site ~200 bp from the CACC at rs17114036 were used as a negative control. n=4. **(D)** Increased enhancer activity of chr1:56962213-56963412 by KLF2 overexpression. Dual-luciferase reporter assays were conducted in HAEC transfected with luciferase constructs containing the PLPP3 promoter and enhancer with either the major (T) or minor (C) allele at s17114036 along with KLF2-overexpressing or control plasmids. KLF2 overexpression resulted in a 2.9-fold increased luciferase activity in HAEC transfected with T allele-containing construct and a 4.7-fold increase in cells transfected with C allele-containing construct. n=3. *p<0.05; **p<0.005; ***p<0.0005 as determined by Student’s t-test. Data represent mean ± SEM. **(E)** The interplay between hemodynamic forces, chromatin landscapes at PLPP3 intron 5, and rs17114036 at the molecular level in regulating endothelial PLPP3 expression and vascular functions.

## Discussion

Although it is proposed that genetic and environmental factors jointly influence the risk of most common human diseases, the interplay between genetic predisposition and biomechanical cues at the molecular level is poorly understood. The biology underlying the majority of CAD and IS GWAS loci remains to be elucidated (28). Most of the CAD and IS SNPs reside in the noncoding genome. Gupta et al. recently reported that the non-coding common variant at rs9349379, implicated in CAD by GWAS, regulates endothelin 1 (EDN1) expression in endothelium (29). Atherosclerotic lesions preferentially develop at elastic arteries where vascular endothelial cells are activated by local disturbed flow (2–6). As of now, it remains unknown whether disease-associated genetic variants contribute to mechano-sensing mechanisms by which cells sense and convert biomechanical stimuli to biological signaling. Our results here elucidate the convergence of CAD/IS genetic predisposition and mechano-transduction mechanisms in endothelial PLPP3 expression. Statistical analyses, whole-genome chromatin accessibility/enhancer marks, genome editing, enhancer assays, chromatin accessibility QTL (caQTL) mapping, and allelic imbalance (AI) assay collectively demonstrate that CAD/IS locus 1p32.2 harbors a mechano-sensitive endothelial enhancer that regulates PLPP3 expression. Moreover, CAD/IS protective allele at rs17114036 confers an increased enhancer activity that is dynamically regulated by unidirectional flow and transcription factor KLF2 (Fig. 5E).

Dysregulation of mechano-sensing mechanisms contributes to the etiology of a wide range of human diseases in cardiovascular, pulmonary, orthopedic, muscular, and reproductive systems (1). The genetic basis of these complex human diseases has been strongly suggested by GWAS but the interplay between genetic variants and mechano-sensing mechanisms has not been investigated. Our data provide the first line of evidence supporting the genetic regulation of mechano-transduction mechanisms in complex human diseases and suggest an underappreciated role of genetic predisposition in cellular mechano-sensing processes.

Transcriptional enhancers orchestrate the majority of cell-type-specific patterns of gene expression (8) and play key roles in development, evolution, and disease (30) which are tightly regulated by mechanical cues (1). Our data provide new molecular evidence that the non-coding genome actively participates in cellular mechanotransduction mechanisms that are influenced by human genetic variances. In addition to the flow-regulation of the specific locus 1p32.2, our results provide one of the first datasets to systematically determine the mechanosensitive chromatin accessibility and putative enhancer regions at the whole-genome scale in vascular endothelium. It is important to note that most of the epigenome studies including ENCODE were conducted in cells without physiological or pathophysiological mechanical stimuli, such as HUVEC under static (no flow) conditions (7). Since major endothelial functions are tightly and dynamically regulated by hemodynamics (2–4), our whole-genome epigenome profiling in HAECs under athero-relevant flows may benefit future studies to investigate mechanical regulation of the non-coding genome in vascular cells. Indeed, we have applied Model-based Analysis of ChIP-Seq (MACS2) (31) and HOMER differential analysis (32) which unbiasedly identified rs17114036-containing locus as one of the 36,965 open chromatin sites that are activated by unidirectional flow (Supplementary Fig. 8).

Mechano-sensitive transcription factors have been proposed as major regulators to determine endothelial functions relevant to atherogenesis. For instance, nuclear factor-κB and HIF-1α mediate gene sets associated with pro-inflammatory, pro-coagulant, and glycolytic endothelial phenotypes under disturbed flow while Krüppel-like factors and nuclear factor erythroid 2–like 2 regulate gene networks promoting the quiescent endothelial phenotype under unidirectional flow (26, 27, 33-36). However, the interaction between flow-sensitive transcription factors and disease-associated genetic predisposition in vascular functions has not been suggested. Our results here demonstrate that a genetic variant can influence important endothelial functions via a non-coding enhancer region recognized by the mechano-sensitive transcription factor KLF2. These results are consistent with emerging evidence showing top-scoring disease-associated SNPs are frequently located within enhancers explicitly active in disease-relevant cell types (10). Moreover, the data suggest that disease-associated genetic variants, via modulation of transcription factor binding, may regulate the enhancer activities dynamically responding to biomechanical cues that are instrumental to key cellular processes.

GWAS related to atherosclerotic diseases have suggested previously unsuspected loci, genes, and biology involved in lipoprotein metabolism (28), resulting in the development of new cholesterol-lowering therapies (37). Despite that dyslipidemia is a major systemic risk factor of CAD and ischemic stroke, atherosclerotic lesions largely initiate and develop at arterial regions of atypical vascular geometry associated with disturbed flow. Previous studies demonstrated that cellular mechano-transduction mechanisms, particularly endothelial responses to hemodynamics, causatively contribute to the focal nature of atherosclerotic lesions (2–6). Our studies here demonstrate that genetic variants not only contribute to inter-individual variation in plasma lipid concentrations (38) but also endothelial responses to blood flow. Indeed, genetic variants at rs17114036 predict CAD susceptibility independent of traditional systemic risk factors such as cholesterol and diabetes mellitus (11, 12). Recent GWAS identified 15 new CAD risk loci near genes of key functions in endothelial, smooth muscle, and white blood cells (39), further highlighting the potential importance of genetic contribution to the arterial-wall-specific mechanisms in atherogenesis. Our results indicate that CAD genetic predisposition and disturbed flow converge to inhibit endothelial PLPP3 expression and restoration of endothelial PLPP3 in atherosusceptible regions may provide an attractive approach for future arterial wall-based atherosclerosis therapy complementary to current pharmacological treatments targeting systemic risk factors.

Our studies demonstrate that the latest human genetics approaches such as caQTL mapping (23), allelic imbalance assay (24), and CRISPR-based assays (20, 21) are powerful tools to investigate possible genetic contributions to cellular mechanotransduction. Miao et al. recently applied CRISPR-Cas9 to achieve high efficiency of a 10 kb deletion of an enhancer region in bulk HUVEC (40). In this study we expanded the applications of CRISPR-based techniques to investigate key vascular functions. Isogenic adult aortic endothelial lines subjected to CRISPR-based deletion were successfully selected to determine the causal role of ~66bp genomic region in regulating endothelial PLPP3 expression. Nevertheless, one limitation here is that we are still unable to replace this human SNP at rs17114036 in adult aortic endothelium even though we have tried various methods to promote homology directed repair during CRISPR-Cas9 gene editing (21). This will be the subject of a future study. Nevertheless, caQTL mapping (23) and allelic imbalance assay (24) provide complementary approaches detecting at the single nucleotide resolution that CAD/IS protective allele at rs17114036 confers a higher enhancer activity at the PLPP3 intron 5. This study demonstrated that human haplotypes and related *cis*-regulatory elements provide an important layer of molecular controls by which cells convert physical stimuli into biological signaling.

## Methods

### Assay for Transposase-Accessible Chromatin using sequencing (ATAC-seq)

ATAC-seq was performed as previously described (18) using Tn5 transposase (Illumina, San Diego CA). Libraries were sequenced on an Illumina HiSeq 4000 according to manufacturer’s specifications by the Genomics Core Facility at the University of Chicago. The reads were aligned to the UCSC hg19 genome using Bowtie2 (41). ATAC-seq were conducted in HAECs under static conditions or subjected to 24-hr unidirectional flow or disturbed flow.

### Chromatin accessibility quantitative trait locus (caQTL) mapping and Allelic Imbalance

caQTL mapping was performed to test for association between genotype at rs17114036 and chromatin accessibility measured by ATAC-seq. We pulled genotypes for HAEC donors from our previous study (42) and imputed linked alleles using IMPUTE2 and SHAPEIT as we published previously (43). Association testing between ATAC-seq tags at the rs17114036 enhancer and genotype were performed using the Combined Haplotype Test in WASP (24).

To perform allelic imbalance (AI) analysis that assigns next generation sequencing reads overlapping heterozygous sites to one chromosome or the other, we quantified ATAC-seq tags at the rs17114036 enhancer using HOMER’s annotatePeaks function to express the log2 normalized tags in this region.

### CRISPR Cas9-mediated deletion of enhancer in teloHAECs

The CRISPR reagents were adapted from the Alt-R system from IDT (IDT, Coralville, IA). The guide RNAs were designed using an online tool at http://crispr.mit.edu/ to minimize off targeting effects using two guides to create a ~66 bp deletion. The guide RNAs were made by annealing the tracrRNA to the sgRNA. Cas9-guide RNA ribonucleoprotein (RNP) complex by incubating guide RNAs with recombinant S. pyogenes Cas9, followed by the delivery to cells using Lipofectamine RNAiMAX (Thermo). For each successive treatment the reagent amounts were scaled relative to the size of the destination vessel to compensate for the number of cells in the reaction. The volumes for each part of the reaction was increased 4x when treating cells from the 96-well to a 6-well, and 16x when moving from the 6-well to the T-75 flask.

Detailed methods are available in the supplemental materials.

---

**Supplemental Table 1.**
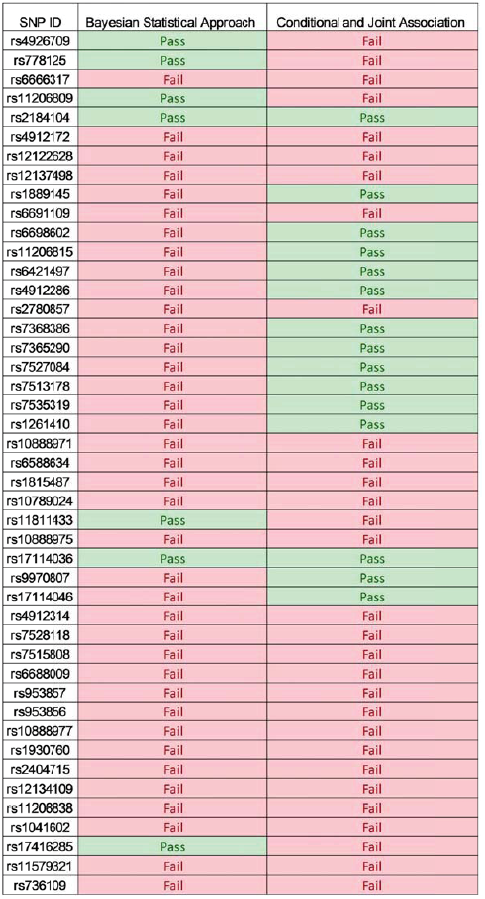
Results from the bioinformatic screening for predicted, causal variants. The results of each test are either “pass” or “fail” highlighted in green or red respectively for each SNP.

**Supplemental Figure 1.**
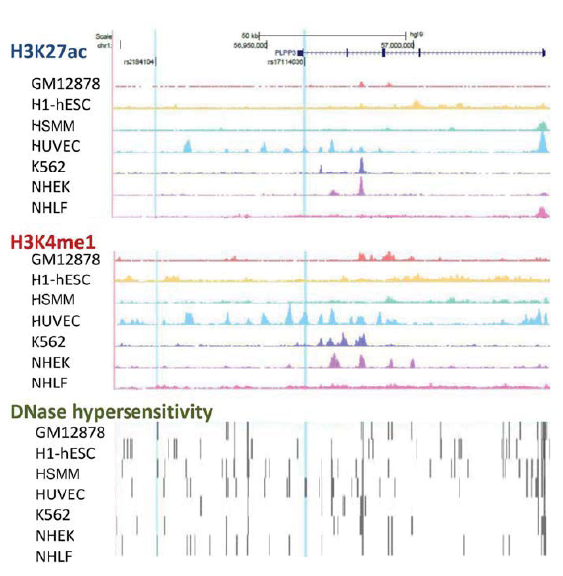
Encyclopedia of DNA Elements (ENCODE) studies indicate rs17114036 is located in an enhancer-like element in Human Umbilical Vein Endothelial Cells (HUVEC). H3K27ac ChIP-seq, H3K4me1 ChIP-seq, DNase hypersensitivity data sets curated by the ENCODE project in GM12878, h1-ESC, HSMM, HUVEC, K562, NHEK, and NHLF cells are plotted at the PLPP3 locus alongside the predicted, causative CAD SNPs rs17114036 and rs2184104.

**Supplemental Figure 2.**
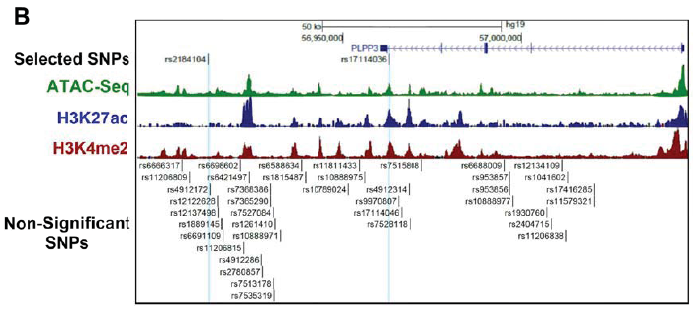
Chromatin accessibility and canonical enhancer marks along with common SNPs in CAD locus 1p32.2 in human aortic endothelial cells. All common SNPs in CAD locus chr 1p32.2 tested for predicted causality are plotted next to the ATAC-seq, H3K27ac ChIP-seq, and H3K4me2 ChIP-seq data performed in HAEC.

**Supplemental Figure 3.**
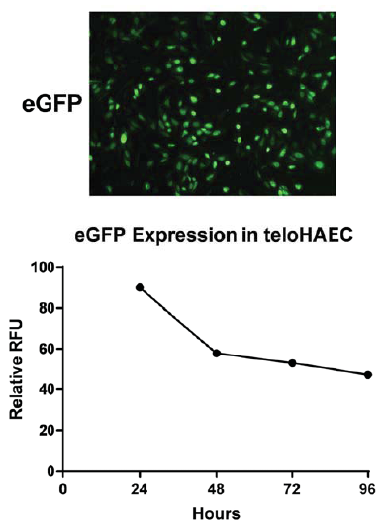
Electroporation results in high transfection efficiency of plasmids in teloHAEC. teloHAEC were transfected with 1.2 μg of eGFP plasmid via electroporation Neon(®) Transfection System (Thermofisher). The cells were visualized under a fluorescent microscope 24 hours post electroporation. GFP measurements were recorded at 24, 48, 72, and 96 hours post-transfection.

**Supplemental Figure 4.**
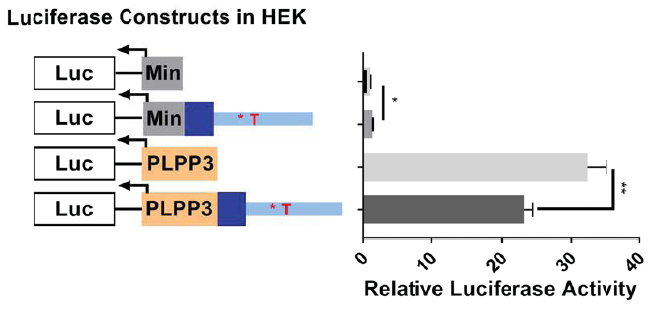
Dual-luciferase reporter assays demonstrate minimal enhancer activity of chr1:56962213-56963412 in embryonic kidney cells 293 (HEK-293). Luciferase reporters constructs with and without the enhancer, as well as with minimal promoter or human PLPP3 promoter were transfected into HEK-293 using lipofectamine and incubated 24 hours before measuring luciferase activity. n=5. **p<0.005

**Supplemental Figure 5.**
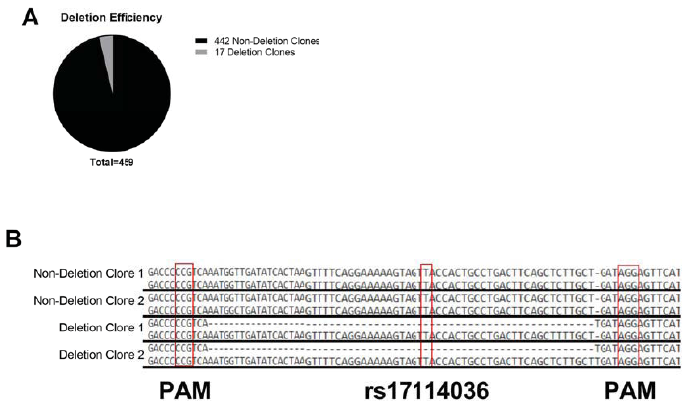
Bacterial CRISPR-associated protein-9 nuclease (CRISPR/Cas9) system results in successful genomic deletion at site of interest in teloHAEC. **(A)** The total numbers of teloHAEC clones that were grown and genotyped are reported. **(B)** The PCR products were cloned (TA cloning) and sequenced to confirm deletion of the targeted region surrounding rs17114036. Protospacer adjacent motif (PAM) sequences, which are recognized and cleaved by Cas9, and rs17114036 are highlighted for reference.

**Supplemental Figure 6.**
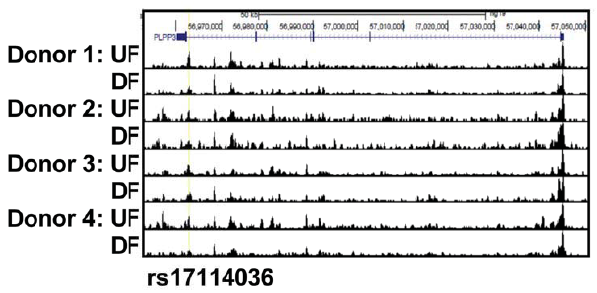
Unidirectional flow (UF) increases chromatin accessibility in PLPP3 intron 5 in HAEC when compared with cells under disturbed flow (DF). ATAC-seq were conducted in HAEC lines from 4 individual donors heterozygous (T/C) at rs17114036 under 24-hr athero-protective UF and athero-susceptible DF.

**Supplemental Figure 7.**
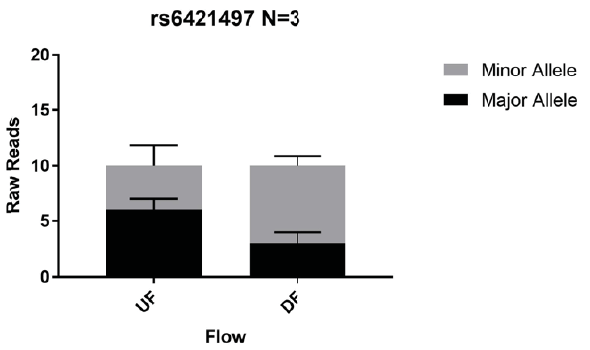
Allelic imbalance analyses show unidirectional flow does not increase chromatin accessibility at locus related to the minor (CAD protective) allele at rs6421497 in human aortic endothelium. SNP rs6421497 was not predicted to be causal but resides within a peak from the ATAC-seq data set in at least 3 donors. The number of ATAC-seq reads detected under unidirectional and disturbed flow are reported and the proportion harboring either the major or minor allele.

**Supplemental Figure 8.**
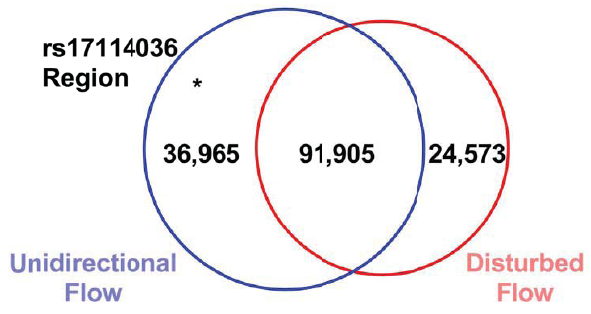
Numbers of open chromatin loci in HAEC regulated by athero-protective unidirectional flow or athero-susceptible disturbed flow. Model-based Analysis of ChIP-Seq (MACS) and HOMER differential analysis were conducted using ATAC-seq results from total 5 HAEC lines subjected to 24 hrs athero-relevant flows to identify hemodynamics-regulated open chromatin sites.

## Supplemental Methods

### Cell culture

Human Aortic Endothelial Cells (HAECs) were procured from Lonza (Allendale, NJ) (CC-2535). HAEC lines that are heterozygous at rs17114036 were from aortic explants of heart transplant donors in the UCLA transplant program and cultured as described previously (1). For CRISPR/Cas9-mediated genome editing, we used teloHAECs (ATCC #CRL-4052) that express hTERT and large-T antigen. teloHAECs were used due to their ability to form colonies, which was necessary for generating isogenic cell lines with deletions. Cells were grown in EGM-2 medium supplemented with SingleQuots from Lonza (CC-3156 & CC-4176) and Antibiotic-Antimycotic from Gibco (Grand Island, NY) (15240062) in a 37°C incubator with 5% CO_2_. HAECs were used from passages 6-10. teloHAECs, and CRISPR-edited teloHAECs were used from passages 10-15. THP-1 cells were maintained using RPMI 1640 medium (Gibco) containing 10% FBS (Biowest).

### H3K27ac and H3K4me2 chromatin immuno-precipitation with whole genome sequencing (ChIP-seq)

HAECs were washed three times with warm PBS and then trypsinized. Cells were pelleted at 3000 x g for 5 min before being fixed at room temperature with 1% paraformaldehyde in PBS for 10 min, and quenched with 125 mM glycine. 1 million cells were used for each ChIP-seq. Cell lysates were sonicated using BioRuptor Pico (Diagenode, Belgium), and then immunoprecipitated using antibodies against H3K27ac (Active Motif, Carlsbad, CA, #39135) or H3K4me2 (EMD Millipore, Billerica, MA, #07– 030), bound to a 2:1 mixture of Protein A Dynabeads (Invitrogen #10002D) and Protein G Dynabeads (Invitrogen #10004D). Following immunopreciptation, crosslinking was reversed and libraries were prepared using the same method described previously (2) for RNA-seq beginning with dsDNA end repair and excluding UDG. For each sample condition, an input library was also created using an aliquot of sonicated cell lysate that had not undergone immunoprecipitation. These samples were sequenced on an Illumina HiSeq 4000 and used to normalize ChIP-seq results.

### RNA-seq

Total RNA was isolated from HAECs using the Quick-RNA Micro Prep kit from ZymoResearch (#R1051), including optional DNase I treatment. mRNA was selected through poly-A isolation using Oligo d(T)25 beads (New England BioLabs #S1419S), fragmented, and cDNA was synthesized with the SuperScript III First-Strand Synthesis System (Invitrogen # 18080051) Second strand synthesis was performed using DNA Polymerase I (Enzymatics #P7050L). To generate sequencing libraries, DNA ends were repaired with T4 DNA Polymerase (Enzymatics #P7080L). Six-mer barcoded adapters (BIOO Scientific NEXTflex #514104) were ligated with T4 DNA Ligase (Enzymatics #L-6030-HC- L) and samples were treated with Uracil DNA Glycosylase (UDG) (Enzymatics #G5010L). Libraries were then amplified less than 14 cycles by PCR (Phusion Hot Start II #F549L) and purified (Zymo #D5205) for high-throughput sequencing at the University of Chicago.

### Normalization of High-throughput Sequencing Data

Reads from ATAC-seq and ChIP-seq were mapped to the hg19 build of the human genome with Bowtie2 (3) and RNA-seq reads were mapped with STAR (4). For RNA-seq, ATAC-seq, and ChIP-seq mapped reads were normalized to 10 million tags per experiment and PCR duplicates were removed in HOMER (5). The ATAC-seq and ChIP-seq aligned files were then converted to BAM format and sorted using SAMtools (6). The files were then filtered to remove unmapped and mitochondrial reads using SAMtools (6). Peak calling was performed using MACS2 (7). The bedgraph files from MACS2 were converted into BigWig files using HOMER for visualization on the genome browser (5). The images in this manuscript had the peaks smoothed to 3 pixels for aesthetics. Using the RNA-seq datasets, PLPP3 expression was quantified using the ‘analyzeRepeats’ function, and is expressed as log2 normalized tag counts. Sequencing data generated in this study is available under Gene Expression Omnibus #GSE112340 (https://www.ncbi.nlm.nih.gov/geo/query/acc.cgi?acc=GSE112340; token: srqrygwoxhajbgx).

### Dual luciferase assay

HAECs and teloHAECs were electroporated using the Neon system (ThermoFisher Scientific). Briefly, the cells were re-suspended in resuspension buffer to have around 40,000 cells per μl. 1.2 μg of eGFP expression plasmid (pmaxGFP) was first used to assess transfection efficiency. For dual luciferase reporter assays, 1 μg of pGL4.23 plasmids carrying firefly luciferase inserted with chr1:56962213-56963412 (T or C at rs17114036, UCSC VERSION hg19) and PLPP3 promoter or minimal promoter at 1 μg/μl and 200 ng of pRL-TK (control reporters) at 1 μg/μl were added per 8.8 μl of cells. The cells were electroporated at 1200 V, for 1 pulse and then were immediately plated onto 12-well culture plates. Media was changed after 4-6 hours to remove dead cells. The peak expression of plasmids was shown at 24 hours post-electroporation and was chosen as the end-point for most luciferase experiments. Cells were collected by adding passive lysis buffer and then put at ‐80°C overnight. Luminescence was measured by Dual-Luciferase Reporter Assay System (Promega) according to manufacturer’s instructions using a Cytation3 plate reader (BioTek).

### CRISPR Cas9-mediated deletion of enhancer in teloHAECs

After four treatments with the CRISPR RNP, treated cells were trypsinized to a single cell suspension. Using a FACS AriaII flow cytometer single cells were plated onto 96 well plates containing EGM-2 media with 10% FBS for a total of 576 possible clones to screen. The cells were incubated for 20 days prior to grow to confluence. The cells were then split and half of each clone was placed onto two new plates to continue growing clones, and to collect DNA for screening. To confirm the CRISPR Cas9-mediated deletion, DNA for screening was collected using 50 μl QuickExtract solution (Epicentre, Madison WI). Mutations were detected using SYBR green (Roche, Indianapolis, IN) and CRISPR Genotyping Screening primers flanking the deletion region. Positive clones were checked with another PCR using Platinum Taq and CRISPR Genotyping TA Cloning primers flanking the deletion. The PCR products for clones were purified using Qiagen PCR clean-up kit (Qiagen, Venlo, Netherlands). The PCR products were ligated into the pGEMT-easy linear vector, cloned, and then submitted for Sanger sequencing to confirm deletions of the target. (CRISPR Genotyping Screening F: 5’- TCCTCCACGTTTAGTTGCCA −3’, R: 5’- AAGGAATCCAGGGTGTAACCG −3’) (CRISPR Genotyping TA Cloning, F: 5’- GGACGCTGGGAATGAGTGAT-3’, R: 5’- ATTGCCCATATCTGCAACCC-3’)

### Leukocyte adhesion assay

Monolayers of teloHAECs were activated using lysophosphatidic acid (LPA) for 3 hours. THP-1 cells at 5-fold number of teloHAECs were pelleted and resuspended in serum-free RPMI and then labeled with 5 M Calcein AM dye (ThermoFisher) for 30 minutes at 37 °C. The cells were then pelleted and resuspended in serum-free RPMI. Labeled THP-1 cells were then added to the teloHAECs and incubated at 37 °C for 1 hour, with gentle rocking every 30 minutes. The cells were then washed extensively 5 times with warm DPBS. Fluorescence of the Calcien AM dye was then measured on a Cytation 3 (Biotek, Winooski, VT) device in area scanning mode, with gain of 80, and excitation at 492 nm, and emission 550 nm.

### Measurement of Transendothelial Electrical Resistance

Cell permeability was evaluated by measuring transendothelial electrical resistance (TER) across CRISPR/Cas9-edited clones on 8 well electrode arrays 8W10E+ (Applied Biophysics) by an electrical cell-substrate impedance sensing system Model 1600R (Applied Biophysics).

### Chromatin Immunoprecipitation PCR

HAECs were transfected with *in vitro* transcripts of KLF2 with HA tag (8) for 6 hours before cross-linking, digestion, and immunoprecipitation according to the Pierce Agarose ChIP Kit. Briefly, cells were cross-linked with 1% formaldehyde in growth media (EGM2). After a 10-minute incubation, glycine was added to 125 mM final concentration and, after 5 minutes, the solution was aspirated. The cells were washed with ice-cold PBS twice, before being scraped in 1mL PBS and 10 μL protease inhibitors (Halt Cocktail, Pierce). The cells were pelleted at 3000 x g for 5 min before undergoing MNase digestion at 10 U/μL for 15 min. After recovery of digested chromatin, the solution was immunoprecipitated with anti-HA antibody at 3 μg/ 2 million cells, and with normal rabbit IgG as control overnight. After binding to agarose beads, the immunoprecipitate was eluted before DNA clean up and then purified DNA was detected by qPCR with PLPP3 enhancer (F: 5’-AGACTAAGACGACGCTCTCC-3’, R: 5’- GTGGCACCTACATCATGTTGT −3’) and PLPP3 promoter primers (F: 5’-TTGCTAACCCTCACAGAGCA-3’, R: 5’-ATCCTGTGACTCTGTGCCTC-3’) as well as negative control primers (F: 5’- ATGTGGCCAGAGTGAAACCA-3’, R: 5’-TCTACACCCAACAGCCTTCT-3’).

### mRNA quantitative real-time PCR

Total RNA was isolated using Direct-zol RNA MiniPrep (Zymo Research). mRNA was reverse-transcribed into cDNA using High Capacity cDNA Reverse Transcription kit (Life Technologies). cDNA quantification was performed on LightCycler 480 II (Roche) using SYBR Green I Master for mRNA. Absolute quantification of each gene of interest was normalized to the geometric mean of beta-actin, ubiquitin B, and GAPDH. The following primer sequences were used (IDT, Coralville, IA):

PPAP2B, 5’-CAGCGCCATGCAAAACTACA-3’ 5’-AAAACCCTCGGTGGTAAGGC-3’
SELE, 5’-CCGAGCGAGGCTACATGAAT-3’ 5’-GCCACATTGGAGCCTTTTGG-3’
GAPDH 5’- TGCACCACCAACTGCTTAGC-3’ 5’- GGCATGGACTGTGGTCATGAG-3’
ACTB 5’- TCCCTGGAGAAGAGCTACGA-3’ 5’- AGGAAGGAAGGCTGGAAGAG-3’
UBB 5’- ATTTAGGGGCGGTTGGCTTT-3’ 5’- TGCATTTTGACCTGTTAGCGG-3’

### Reagents and antibodies

Anti-HA antibody was from Abcam (ab9110). Lysophosphatidic acid (LPA) was obtained from Santa Cruz Biotechnology (Cat# 22556-62-3). High molecular weight dextran was obtained from Sigma (Cat# 31392-50G).

### Application of athero-relevant flows

Hemodynamic forces were applied to cultured HAECs in a 6 well plate using a 1° tapered stainless steel cone held by a computerized stepper motor UMD-17 (Arcus Technology, Livermore CA). The cone rotation recapitulated the shear stress waveform mimicking the disturbed flow at the athero-susceptible human carotid artery or the shear stress profile mimicking the unidirectional flow at the athero-resistant distal internal carotid artery (9). The flow device was placed in a 37°C incubator with 5% CO2. HAECs at 100% confluence, maintained in EGM2- medium containing 4% dextran (Sigma-Aldrich, St. Louis, MO, 31392) in 6-well plates, were subjected to unidirectional flow or disturbed flow for 24 hours before being processed.

## References

1. Jaalouk DE & Lammerding J (2009) Mechanotransduction gone awry. Nat Rev Mol Cell Biol 10(1):63–73.

2. Hahn C & Schwartz MA (2009) Mechanotransduction in vascular physiology and atherogenesis. Nat Rev Mol Cell Biol 10(1):53–62.

3. Gimbrone MA, Jr. & Garcia-Cardena G (2016) Endothelial Cell Dysfunction and the Pathobiology of Atherosclerosis. Circ Res 118(4):620–636.

4. Davies PF (2009) Hemodynamic shear stress and the endothelium in cardiovascular pathophysiology. Nat Clin Pract Cardiovasc Med 6(1):16–26.

5. Kumar S, Kim CW, Simmons RD, & Jo H (2014) Role of flow-sensitive microRNAs in endothelial dysfunction and atherosclerosis: mechanosensitive athero-miRs. Arterioscler Thromb Vasc Biol 34(10):2206–2216.

6. Shyy JY & Chien S (2002) Role of integrins in endothelial mechanosensing of shear stress. Circ Res 91(9):769–775.

7. Consortium EP (2012) An integrated encyclopedia of DNA elements in the human genome. Nature 489(7414):57–74.

8. Ong CT & Corces VG (2011) Enhancer function: new insights into the regulation of tissue-specific gene expression. Nat Rev Genet 12(4):283–293.

9. Heinz S, Romanoski CE, Benner C, & Glass CK (2015) The selection and function of cell type-specific enhancers. Nat Rev Mol Cell Biol 16(3):144–154.

10. Ernst J, et al. (2011) Mapping and analysis of chromatin state dynamics in nine human cell types. Nature 473(7345):43–49.

11. Schunkert H, et al. (2011) Large-scale association analysis identifies 13 new susceptibility loci for coronary artery disease. Nat Genet 43(4):333–338.

12. Deloukas P, et al. (2013) Large-scale association analysis identifies new risk loci for coronary artery disease. Nat Genet 45(1):25–33.

13. Dichgans M, et al. (2014) Shared genetic susceptibility to ischemic stroke and coronary artery disease: a genome-wide analysis of common variants. Stroke 45(1):24–36.

14. Wu C, et al. (2015) Mechanosensitive PPAP2B Regulates Endothelial Responses to Atherorelevant Hemodynamic Forces. Circ Res 117(4):e41–53.

15. Panchatcharam M, et al. (2014) Mice with targeted inactivation of ppap2b in endothelial and hematopoietic cells display enhanced vascular inflammation and permeability. Arterioscler Thromb Vasc Biol 34(4):837–845.

16. Maller JB, et al. (2012) Bayesian refinement of association signals for 14 loci in 3 common diseases. Nat Genet 44(12):1294–1301.

17. Yang J, et al. (2012) Conditional and joint multiple-SNP analysis of GWAS summary statistics identifies additional variants influencing complex traits. Nat Genet 44(4):369–375, S361–363.

18. Buenrostro JD, Giresi PG, Zaba LC, Chang HY, & Greenleaf WJ (2013) Transposition of native chromatin for fast and sensitive epigenomic profiling of open chromatin, DNA-binding proteins and nucleosome position. Nat Methods 10(12):1213–1218.

19. Abecasis GR, et al. (2012) An integrated map of genetic variation from 1,092 human genomes. Nature 491(7422):56–65.

20. Ran FA, et al. (2013) Genome engineering using the CRISPR-Cas9 system. Nat Protoc 8(11):2281–2308.

21. Lin S, Staahl BT, Alla RK, & Doudna JA (2014) Enhanced homology-directed human genome engineering by controlled timing of CRISPR/Cas9 delivery. Elife 3:e04766.

22. Dai G, et al. (2004) Distinct endothelial phenotypes evoked by arterial waveforms derived from atherosclerosis-susceptible and ‐resistant regions of human vasculature. Proc Natl Acad Sci U S A 101(41):14871–14876.

23. Kumasaka N, Knights AJ, & Gaffney DJ (2016) Fine-mapping cellular QTLs with RASQUAL and ATAC-seq. Nat Genet 48(2):206–213.

24. van de Geijn B, McVicker G, Gilad Y, & Pritchard JK (2015) WASP: allele-specific software for robust molecular quantitative trait locus discovery. Nat Methods 12(11):1061–1063.

25. Dekker RJ, et al. (2006) KLF2 provokes a gene expression pattern that establishes functional quiescent differentiation of the endothelium. Blood 107(11):4354–4363.

26. SenBanerjee S, et al. (2004) KLF2 is a novel transcriptional regulator of endothelial proinflammatory activation. J Exp Med 199(10):1305–1315.

27. Parmar KM, et al. (2006) Integration of flow-dependent endothelial phenotypes by Kruppel-like factor 2. J Clin Invest 116(1):49–58.

28. Nurnberg ST, et al. (2016) From Loci to Biology: Functional Genomics of Genome-Wide Association for Coronary Disease. Circ Res 118(4):586–606.

29. Gupta RM, et al. (2017) A Genetic Variant Associated with Five Vascular Diseases Is a Distal Regulator of Endothelin-1 Gene Expression. Cell 170(3):522–533 e515.

30. Shlyueva D, Stampfel G, & Stark A (2014) Transcriptional enhancers: from properties to genome-wide predictions. Nat Rev Genet 15(4):272–286.

31. Zhang Y, et al. (2008) Model-based analysis of ChIP-Seq (MACS). Genome Biol 9(9):R137.

32. Heinz S, et al. (2010) Simple combinations of lineage-determining transcription factors prime cis-regulatory elements required for macrophage and B cell identities. Mol Cell 38(4):576–589.

33. Dai G, et al. (2007) Biomechanical forces in atherosclerosis-resistant vascular regions regulate endothelial redox balance via phosphoinositol 3-kinase/Aktdependent activation of Nrf2. Circ Res 101(7):723–733.

34. Hamik A, et al. (2007) Kruppel-like factor 4 regulates endothelial inflammation. J Biol Chem 282(18):13769–13779.

35. Wu D, et al. (2017) HIF-1alpha is required for disturbed flow-induced metabolic reprogramming in human and porcine vascular endothelium. Elife 6:e25217.

36. Ajami NE, et al. (2017) Systems biology analysis of longitudinal functional response of endothelial cells to shear stress. Proc Natl Acad Sci U S A 114(41):10990–10995.

37. Sabatine MS, et al. (2017) Evolocumab and Clinical Outcomes in Patients with Cardiovascular Disease. N Engl J Med 376(18):1713–1722.

38. Willer CJ, et al. (2013) Discovery and refinement of loci associated with lipid levels. Nat Genet 45(11):1274–1283.

39. Howson JMM, et al. (2017) Fifteen new risk loci for coronary artery disease highlight arterial-wall-specific mechanisms. Nat Genet 49(7):1113–1119.

40. Miao Y, et al. (2018) Enhancer-associated long non-coding RNA LEENE regulates endothelial nitric oxide synthase and endothelial function. Nat Commun 9(1):292.

41. Langmead B & Salzberg SL (2012) Fast gapped-read alignment with Bowtie 2. Nat Methods 9(4):357–359.

42. Romanoski CE, et al. (2010) Systems genetics analysis of gene-by-environment interactions in human cells. Am J Hum Genet 86(3):399–410.

43. Hogan NT, et al. (2017) Transcriptional networks specifying homeostatic and inflammatory programs of gene expression in human aortic endothelial cells. Elife 6:e22536.

## References for Supplemental Methods

1. Navab M, et al. (1988) Monocyte migration into the subendothelial space of a coculture of adult human aortic endothelial and smooth muscle cells. J Clin Invest 82(6):1853–1863.

2. Hogan NT, et al. (2017) Transcriptional networks specifying homeostatic and inflammatory programs of gene expression in human aortic endothelial cells. Elife 6:e22536.

3. Langmead B & Salzberg SL (2012) Fast gapped-read alignment with Bowtie 2. Nat Methods 9(4):357–359.

4. Dobin A, et al. (2013) STAR: ultrafast universal RNA-seq aligner. Bioinformatics 29(1):15–21.

5. Heinz S, et al. (2010) Simple combinations of lineage-determining transcription factors prime cis-regulatory elements required for macrophage and B cell identities. Mol Cell 38(4):576–589.

6. Li H, et al. (2009) The Sequence Alignment/Map format and SAMtools. Bioinformatics 25(16):2078–2079.

7. Feng J, Liu T, Qin B, Zhang Y, & Liu XS (2012) Identifying ChIP-seq enrichment using MACS. Nat Protoc 7(9):1728–1740.

8. Huang RT, et al. (2017) Experimental Lung Injury Reduces Kruppel-like Factor 2 to Increase Endothelial Permeability via Regulation of RAPGEF3-Rac1 Signaling. Am J Respir Crit Care Med 195(5):639–651.

9. Dai G, et al. (2004) Distinct endothelial phenotypes evoked by arterial waveforms derived from atherosclerosis-susceptible and -resistant regions of human vasculature. Proc Natl Acad Sci U S A 101(41):14871–14876.

